# Gramicidin increases lipid flip-flop in symmetric and asymmetric lipid vesicles

**DOI:** 10.1101/383463

**Authors:** M. Doktorova, F. A. Heberle, D. Marquardt, R. Rusinova, L. Sanford, T. A. Peyear, J. Katsaras, G. W. Feigenson, H. Weinstein, O. S. Andersen

## Abstract

Unlike most transmembrane proteins, phospholipids can migrate from one leaflet of the membrane to the other. Because this spontaneous lipid translocation (flip-flop) tends to be very slow, cells facilitate the process with enzymes that catalyze the transmembrane movement and thereby regulate the transbilayer lipid distribution. Non-enzymatic membrane-spanning proteins with unrelated primary functions have also been found to accelerate lipid flip-flop in a nonspecific manner and by various hypothesized mechanisms. Using deuterated phospholipids, we examined the acceleration of flip-flop by gramicidin channels which have well-defined structures and known function, features that make them ideal candidates for probing the protein-membrane interactions underlying lipid flip-flop. To study compositionally and isotopically asymmetric proteoliposomes containing gramicidin, we expanded a recently developed protocol for the preparation and characterization of lipid-only asymmetric vesicles. Channel incorporation, conformation, and function were examined with small-angle X-ray scattering, circular dichroism and a stopped-flow spectrofluorometric assay, respectively. As a measure of lipid scrambling we used differential scanning calorimetry to monitor the effect of gramicidin on the melting transition temperatures of the two bilayer leaflets. The two calorimetric peaks of the individual leaflets merged into a single peak over time suggestive of scrambling activity, and the effect of the channel on the transbilayer lipid distribution in both symmetric POPC and asymmetric POPC/DMPC vesicles was quantified from proton NMR measurements. Our results show that gramicidin increases lipid flip-flop in a complex, concentration-dependent manner. To determine the molecular mechanism of the process we used molecular dynamics simulations and further computational analysis of the trajectories to estimate the amount of membrane deformation in the samples. Together, the experimental and computational approaches were found to constitute an effective means for studying the effects of transmembrane proteins on lipid distribution in both symmetric and asymmetric model membranes.

## INTRODUCTION

Membranes are an essential component of all living organisms. Their structure and organization serve many functions and are tightly regulated by the cell. One prominent example is the transverse lipid distribution in cell membranes: whereas a self-assembled lipid bilayer has the same lipid composition in its two leaflets (i.e., it is symmetric), the leaflet compositions in the plasma membranes of eukaryotic cells differ (i.e., the bilayer is asymmetric), and this difference is actively maintained by the cell. Not surprisingly, the biophysical mechanisms underlying membrane asymmetry are the subject of intense studies [1–5] which are rapidly increasing in number as a result of recent advances that have enabled the preparation and biophysical characterization of asymmetric lipid-only model membranes in vitro [6–8]. Because such model membranes are not at chemical equilibrium and their asymmetry is not actively maintained, the time window for examining their properties is limited by the gradual redistribution of the lipids between the two leaflets until a symmetric lipid composition is achieved. Such unassisted interleaflet lipid movement is often referred to as *spontaneous lipid flip-flop (SLF)* [9].

The key lipid and bilayer properties that determine the kinetics of SLF include chain length, headgroup size and charge, and cholesterol concentration (see Ref. [10] for a current review of both experimental and computational studies on the topic). In general, the flip-flop rates of phospholipids are many orders of magnitude slower than other typical lipid motions, such as rotation or lateral diffusion [11], due to the large energy barrier for translocating a polar lipid headgroup from one side of the bilayer to the other, a process that requires traversing the bilayer’s hydrophobic core. In cells, the one-directional movement of lipids between the extracellular and intracellular leaflets is catalyzed by ATP-dependent enzymes such as flippases that move phospholipids from the extracellular leaflet to the intracellular leaflet (e.g., P-type ATPases) [12] and floppases that move phospholipids from the intracellular leaflet to the extracellular leaflet (e.g., ABC transporters) [13], respectively. An additional bi-directional, and often non-specific movement of lipids across the two leaflets is catalyzed by ATP-independent scramblases, which include the Ca^2+^ activated TMEM (TransMEMbrane) family of proteins [11]. It is through this careful balance between the activity of the enzymes and scramblases that cells maintain the desired lipid compositions in their two membrane leaflets.

In addition to the flippases, floppases and scramblases, a wide variety of proteins with non-related primary functions can catalyze lipid flip-flop through different proposed mechanisms that include pore-mediated scrambling [14, 15] and the so-called “credit-card” lipid movement [11]. For some proteins this “side” function has been proposed to have biological implications [16]; for others the physiological role, if any, remains unclear. Still, the ability of a variety of membrane proteins to scramble lipids has direct implications for the design and interpretation of studies employing asymmetric protein-laden membranes and therefore needs to be carefully examined. A particularly interesting case are single-span transmembrane (TM) proteins that often are used to study protein-membrane interactions in vitro [17]. Such proteins can facilitate lipid flip-flop through the so-called perturbation-mediated mechanisms, that is, lowering the energy barrier for lipid translocation between the leaflets by changing the bilayer structure and/or dynamics in the vicinity of the protein [18–22].

One single-span TM protein, which reportedly affects lipid flip-flop only under certain conditions, is the functional dimer form of the bacterial ion-channel gramicidin (gA) [23]. To our knowledge, the channel has been shown to accelerate lipid translocation in three separate studies, but to different extents: up to 30-fold in erythrocytes at moderate gA concentrations (gA:lipid ratio of about 1:200) [24]; from 2 to 10-fold in supported lipid bilayers at high gA:lipid ratios (1:50) [25]; and to a somewhat lesser extent in 400 nm diameter liposomes with high gramicidin ratios (1:20) in which flip-flop was unmeasurable in the absence of gA [14]. At the same time, flip-flop enhancement was not detected in erythrocytes at gA concentrations at which the channel performs its ion-conducting function [24]. The disparate results from these studies highlight limitations in the experimental methodologies, which include quantifying the distribution of lipids in the two leaflets using rather indirect methods, e.g. extraction with albumin (which in its own right could perturb the bilayer) [24], or approximating the kinetics of lipid flip-flop by measuring the translocation rate of a chain-labeled fluorescent lipid analog [14]. The choice of system is also important: complex and asymmetric cell membrane environments like erythrocytes present challenges both for the interpretation of the results and the quantification of the gA:lipid ratio, whereas the unavoidable bilayer defects in supported lipid bilayers prepared with the Langmuir-Blodgett/Langmuir-Schaefer (LB/LS) technique could accelerate lipid flip-flop in a manner that is difficult to control [26].

Here we address these challenges using a new platform for measuring protein-mediated lipid flip-flop in vitro. The approach utilizes free-floating and stress-free large unilamellar vesicles with precisely controlled symmetric and asymmetric lipid composition, both in the presence and absence of protein. This experimental setup allows for a wide range of biophysical assays and, importantly, enables the direct measurement of the flip-flop kinetics of unlabeled lipids in chemically symmetric and asymmetric bilayers using ^1^H NMR. Thus this framework overcomes many of the limitations in the previous approaches by providing a robust model system for studying the effects of proteins on the transverse bilayer organization. Using this protocol, we examined the effect of gramicidin on lipid flip-flop in both isotopically and compositionally asymmetric vesicles. Our results show that gramicidin speeds up lipid translocation in both systems, and that the rates are not directly proportional to gA concentration. Further computational analysis revealed that membrane deformations likely play a role in the observed effects at high gA mole fractions, suggesting the existence of mechanistically different regimes of gA-mediated changes in bilayer organization.

## MATERIALS AND METHODS

### Materials

Gramicidin and all phospholipids (POPC, POPG, POPC-d31, POPC-d13 and DMPC-d54, see Table 1 for a list of abbreviations) were purchased from Avanti Polar Lipids (Alabaster, AL) as dry powders and used as supplied. The phospholipids were dissolved in HPLC-grade chloroform and gramicidin was dissolved in HPLC-grade methanol. Both phospholipids and gramicidin were stored at −80°C until use. Methyl-β-cyclodextrin (MβCD), praseodymium(III) nitrate hexahydrate {Pr(NO3)3 6H2O}, sucrose, NaCl, and hydrochloric acid were purchased from Fisher Scientific. Thallium nitrate (T1NO_3_) and ANTS were purchased from Sigma Aldrich and Invitrogen, respectively. Ultrapure H_2_O was obtained from a High-Q purification system (Wilmette, IL) and D_2_O (99.9%) was purchased from Cambridge Isotopes (Andover, MA).

**Table 1.**
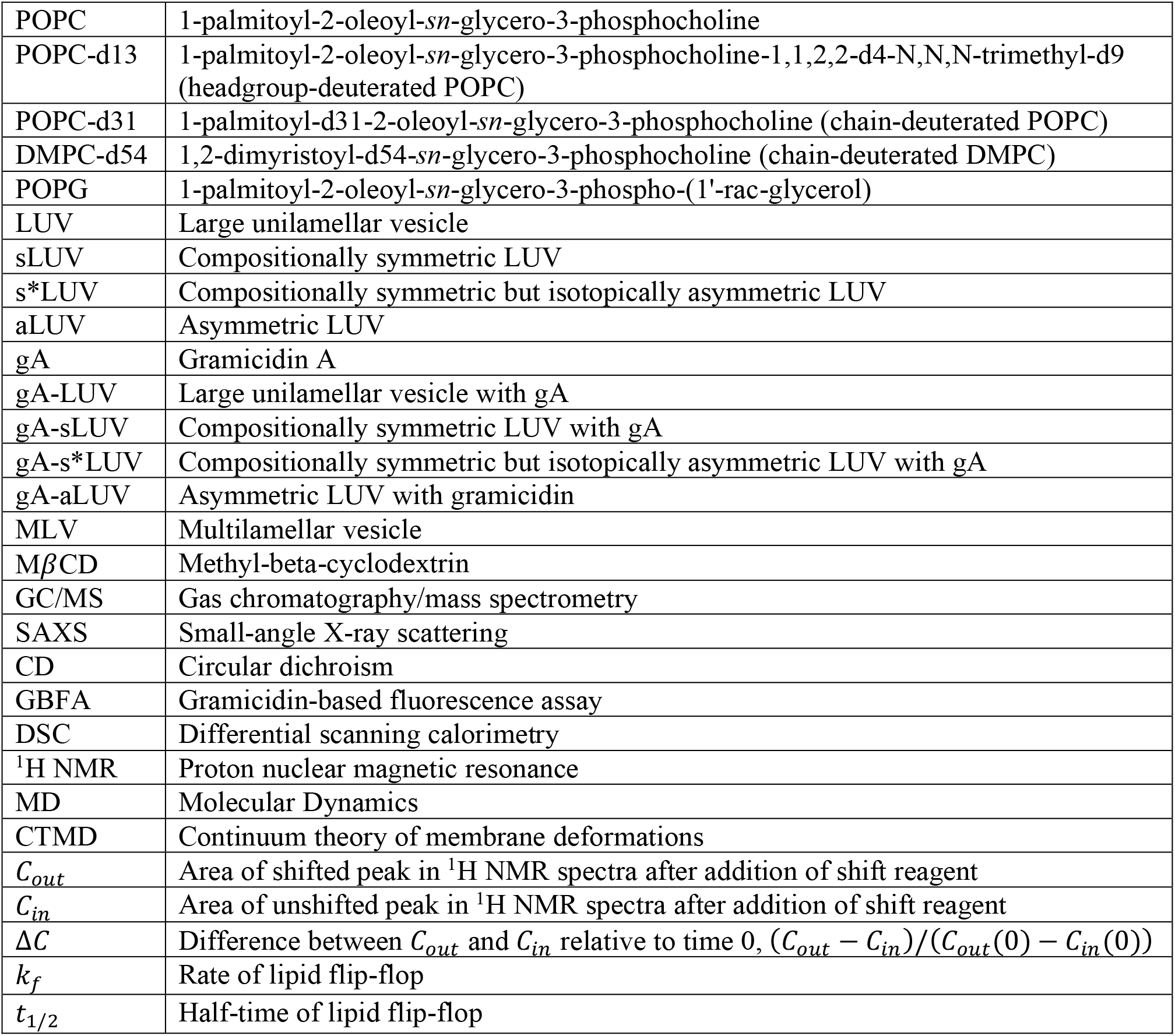
List of abbreviations

### Preparation of liposomes without gramicidin

Large unilamellar vesicles (LUVs) with symmetric lipid distribution (sLUVs) were prepared by first mixing appropriate volumes of lipid stocks in organic solvent with a glass Hamilton syringe. The solvent was evaporated with an Ar stream followed by high vacuum overnight. The dry lipid film was hydrated at room temperature (for POPC) or 35-40°C (for mixtures with DMPC) for at least 1 h with intermittent vortexing every 15 min. The resulting multilamellar vesicle (MLV) suspension was subjected to at least 5 freeze/thaw cycles using a −80°C freezer, then extruded through a 100 nm pore-diameter polycarbonate filter with a mini-extruder (Avanti Polar Lipids) by passing the suspension through the filter 31 times. All sLUVs were doped with 5 mol% POPG to ensure unilamellarity [27].

Asymmetric LUVs (aLUVs) were prepared following the protocol described in [28] with slight modifications. Briefly, an MLV suspension of the donor lipid (i.e., the lipid to be exchanged into the outer leaflet of the aLUVs) was prepared in a 20% w/w sucrose solution as described above. The sample was diluted 20-fold with water and centrifuged at 20°C for 30 min at 20K × g. The supernatant was discarded, the pellet was re-dissolved using a 35 mM MβCD solution in water at a nominal donor lipid:MβCD ratio of 1:8, and the MβCD/donor mixture was incubated at room temperature for 2 h with gentle stirring. sLUVs of the acceptor lipids (i.e., the lipids to be present on the inner leaflet of the aLUVs) were prepared in a 25 mM NaCl solution as described above. These were added to the MβCD/donor mixture at a nominal donor:acceptor ratio of 2:1 or 3:1 (see Table S1). The MβCD/donor/acceptor slurry was incubated for 30 min at room temperature with gentle stirring. Immediately after this incubation the sample was filtered with a centrifugal filter device (Amicon Ultra-15, 100,000 Da molecular weight cutoff), pre-washed with water 7 times) for 25 min at 2.5K × g. The retentate was diluted with water 8-fold and centrifuged for 30 min at 20K × g and 20°C. The supernatant was carefully transferred to centrifugal devices (as described above), concentrated to 250-500 μL and washed with H_2_O or D_2_O a minimum of 3 times. The aLUVs were recovered from the final retentate and stored in a plastic centrifuge tube at room temperature until further use.

### Preparation of liposomes with gramicidin

Symmetric large unilamellar vesicles with gramicidin (gA-sLUVs) were prepared as follows. The lipids were first mixed in organic solvent, and gA was added from a methanolic solution with a glass Hamilton syringe at the appropriate gA:lipid ratio. The organic solvent was evaporated on a rotary evaporator, followed by high vacuum overnight. The dry sample was gently hydrated on a rotary evaporator for 30-60 min, then incubated at room temperature until the film was fully dissolved and the solution looked uniform (typically overnight). Vortexing was performed only occasionally and at the lowest setting. The sample was subjected to at least 5 freeze/thaw cycles using a −80°C freezer and then extruded with a 100 nm pore-diameter polycarbonate filter as described above. All gA-sLUVs were doped with 5 mol% POPG to ensure unilamellarity.

Asymmetric LUVs with gramicidin (gA-aLUVs) were prepared following the protocol described above with the only modification that gramicidin was added to the symmetric acceptor liposomes as described above. That is, instead of acceptor sLUVs, acceptor gA-sLUV vesicles were added to the MβCD/donor mixture after the 2 h incubation. All other steps of the protocol were the same (see Fig. S1).

### Gas chromatography-mass spectrometry (GC/MS)

The overall lipid composition of the aLUVs and gA-aLUVs was determined from GC/MS analysis of fatty acid methyl esters (FAMEs) generated by acid-catalyzed methanolysis. The detailed protocol is described in [28]. Briefly, about 100 μg of the sample were dried on a rotary evaporator, dissolved in 1 mL of 1 M methanolic HCl, flushed with Ar and incubated at 85°C for 1 h in a glass culture tube sealed with a Teflon-lined cap. After allowing the sample to cool for a few minutes, 1 mL water was added and the sample was briefly vortexed. Then, 1 mL hexane was added and the sample was vortexed vigorously to form an emulsion and extract the FAMEs. The solution was centrifuged at low speed (~ 400 × g) for 5 min to break the emulsion, and the upper (hexane) phase containing the FAMEs was transferred to a GC autosampler vial. The total volume in the vial was brought up to 1 mL with hexane. FAME analysis was performed using an Agilent 5890A gas chromatograph (Santa Clara, CA) with a 5975C mass-sensitive detector operating in electron-impact mode and an HP-5MS capillary column (30 m × 0.25 mm, 0.25 μm film thickness). After injection of a 5 uL aliquot into the chromatograph, a pre-programmed column temperature routine was initiated as described in [28]. Total ion chromatogram peaks were assigned and integrated using GC/MSD ChemStation Enhanced Data Analysis software. The ratio of the different lipid components was determined from the ratio of the respective peak areas of the FAMEs corresponding to the lipid fatty acid chains.

### Small-angle X-ray Scattering (SAXS)

LUV samples for small angle X-ray scattering (SAXS) measurements were prepared as described above and concentrated to 15–20 mg/mL. SAXS measurements were performed using a Rigaku BioSAXS-2000 home source system with a Pilatus 100K 2D detector and a HF007 copper rotating anode (Rigaku Americas, The Woodlands, TX). SAXS data were collected at a fixed sample-to-detector distance (SDD) using a silver behenate calibration standard, with a typical data collection time of 3-4 h. The one-dimensional scattering intensity *I*(*q*) [*q* = 4πsin(0)/A, where *λ* is the X-ray wavelength and 2*θ* is the scattering angle relative to the incident beam] was obtained by radial averaging of the corrected 2D image data, an operation that was performed automatically using Rigaku SAXSLab software. Data were collected in 10 min frames with each frame processed separately to assess radiation damage; there were no significant changes in the scattering curves over time. Background scattering from water collected at the same temperature was subtracted from each frame, and the background-corrected *I*(*q*) from the individual frames was then averaged, with the standard deviation taken to be the measurement uncertainty.

### Circular dichroism (CD)

Samples for CD were first diluted to 1 mg/mL lipid concentration, for a protein concentration between 5–13 μM. The lipid concentration was estimated from dynamic light scattering (DLS). gA conformation was measured using a J-815 spectropolarimeter equipped with a PTC-423S Peltier temperature controller (Jasco, Easton, MD). CD spectra (raw ellipticity *θ* in units of millidegree vs. wavelength) were collected at 25°C using a 2 mm path length cuvette, a scan rate of 100 nm/min, and 30 accumulations. The spectrum of each gA-containing sample was first corrected for the lipid background by subtracting the spectrum of a corresponding lipid-only sample. The only exception was the gA-aLUV sample, for which the background was taken to be the spectrum of the POPC acceptor sLUVs, which were similar in size to the gA-aLUVs, as determined from DLS. The background-corrected data were then converted to mean residue molar ellipticity [*θ*] (in units of degree cm^2^ dmol^-1^) using the relationship:

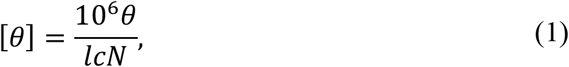

where *l* is the cell path length in mm, *c* is the protein concentration in μM, and *N* = 15 is the number of amino acids in the gA protein.

### Gramicidin-based fluorescence assay (GBFA)

Gramicidin function was quantified using a fluorescence quench protocol [29]. For the present studies the acceptor sLUVs were prepared with a gA:lipid ratio of 1:20,000 and hydrated with 25 mM ANTS instead of 25 mM NaCl. Following the last concentration step, the gA-aLUVs were washed once with H_2_O and 3 times with elution buffer (35 mM NaNO_3_ and 6 mM HEPES at pH 7.0). The rate of quenching of the ANTS trapped inside the vesicles was measured with a Tl+ buffer (35 mM TINO_3_ and 6 mM HEPES at pH 7.0) in a single-mixing experiment and quantified with a regular stretched exponential [29]. As a control, gA-sLUV acceptors hydrated with ANTS were also washed 3 times with elution buffer and the rate of ANTS quenching was measured as described above.

### Differential scanning calorimetry (DSC)

Samples for DSC were diluted to ~ 5 mg/mL and measured using a Nano DSC (TA Instruments, New Castle, DE). The LUV suspension was loaded into the sample capillary cell, and degassed solvent was loaded into the reference capillary cell. The cells were pressurized to 3 atm to suppress the formation of air bubbles, and a cooling scan was initiated from 30°C to −8°C at a rate of 0.2°C/min. All thermograms showed either a series of peaks or a single broad peak between ~ −5°C and 20°C. A sloping background contribution was accounted for by fitting a portion of the thermogram on either side of the peaks of interest to a third-order polynomial which was then subtracted from the raw data.

### ^1^H NMR

The interleaflet lipid distribution in the aLUVs and gA-aLUVs was quantified with a shift reagent assay performed on either a Bruker Avance III 400 MHz spectrometer (at ORNL) or a Bruker Avance III HD 500 MHz spectrometer (at Weill Cornell Medical College) as described in [26]. Briefly, a standard ^1^H pulse sequence with a 30° flip angle and 2 s delay time was collected at 35°C. For all samples measured in H_2_O, the signal from the solvent was suppressed using the standard excitation sculpting sequence *zgesgp*. A sample aliquot was first diluted with D_2_O or H_2_O to a total volume of 600 μL and a concentration of ~ 0.5 mg/mL. The diluted sample was loaded into an NMR tube and a spectrum was recorded. A small aliquot (1-2 μL) of 20 mM Pr^3^+/D_2_O was then added to the NMR tube and mixed with its contents by inverting the tube 3 times ([Pr^3^+] ≈ 50 μM), and spectrum was recorded immediately thereafter; another aliquot of Pr^3+^ was added, and the procedure was repeated. A total of at least three Pr^3+^ titrations were performed, and their spectra were recorded and used to establish the measurement uncertainty as described below. The typical elapsed time between the first exposure of the sample to Pr^3^+ and completion of the experiment was 40 min. Each Pr^3^+ titration resulted in a further shift of the resonance peak of the protiated choline headgroups exposed on the outer leaflet of the vesicles as shown in Fig. S2.

NMR analysis was performed by fitting each spectrum as a sum of Lorentzians using Origin or custom Mathematica scripts. The choline resonances were identified, and the fraction of exposed and protected choline headgroups was quantified from the areas of the shifted and unshifted choline peaks and used to determine the fraction of protiated-headgroup lipid in each leaflet. The measurement uncertainty was calculated as the standard deviation of peak areas determined from the Pr^3+^ titration data described above. This information, together with the GC results, was used to calculate the composition of the two bilayer leaflets as described in [28, 30].

### Analysis of lipid flip-flop kinetics

The rate of lipid flip-flop, *k_f_*, was measured from the time-dependent changes in the transverse distribution of protiated-headgroup lipids in each sample as described in [26]. Briefly, the NMR time traces of relative changes in the lipid distribution were expressed as:

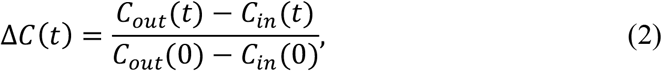

where *C(t)* is the area of the shifted (outer leaflet) and unshifted (inner leaflet) choline peaks at time *t* with subscripts *out* and *in* denoting the outer and inner leaflet, respectively. The data was modeled as:

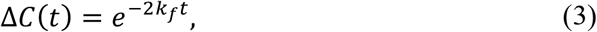

with the corresponding half-time *t_1/2_* given by

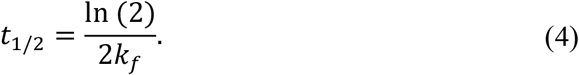

All samples were kept on the bench between NMR measurements, and consequently the flip-flop kinetics reported here correspond to sample behavior at an ambient temperature of ~ 22°C.

### Molecular Dynamics (MD) simulations

All-atom MD simulations of gramicidin in the symmetric and compositionally asymmetric bilayers from the *in vitro* experiments were performed as described below.

For the symmetric system, a POPC bilayer with 100 lipids per leaflet (200 lipids total) was first constructed with CHARMM-GUI [31–33]. The bilayer was hydrated with 70 waters per lipid, and a total of 35 Na+ and 35 Cl^-^ ions for a salt concentration of 138 mM. After a short initial equilibration with CHARMM-GUI’s protocol, the system was run for 226 ns. From the last frame of the trajectory all water and ion atoms were removed, and a single gramicidin dimer (PDB: 1JNO) was manually inserted in the bilayer by replacing 10 lipids from each leaflet with a gA monomer. The system was then hydrated and ionized with VMD’s *solvate* and *autoionize* plugins, respectively, resulting in a bilayer patch with 90 lipids per leaflet (180 lipids total), 1 gA dimer, 67 waters per lipid and 30 Na+ and 30 Cl^−^ ions for a salt concentration of 138 mM NaCl.

For the asymmetric system, we first identified the appropriate number of lipids in the asymmetric lipid-only bilayer with top leaflet composed of DMPC/POPC 75/25 mol%, and a bottom leaflet composed of DMPC/POPC 10/90 mol%, following the protocol in [34]. The resulting tension-free bilayer contained 104 and 100 lipids in the top and bottom leaflets respectively, and was constructed and equilibrated with CHARMM-GUI’s protocols. After a production run of 445 ns, gramicidin was manually inserted in the bilayer by removing 8 lipids from each leaflet while maintaining the overall leaflet compositions. The system was hydrated with VMD’s *solvate* plugin, resulting in a bilayer patch with 96 lipids in the top leaflet, 92 lipids in the bottom leaflet, 1 gA dimer and 55 waters per lipid.

All simulations were performed with the NAMD software [35] and the CHARMM36 force field for lipids [36, 37] in the NPT ensemble under constant pressure of 1 atm and temperature of 25°C. The force field parameters for gA were kindly provided by Andrew Beaven and were based on those used in [38], made compatible with the CHARMM36 force field. Namely, the D-amino acids were treated in the same way as L-amino acids (except for their chirality) whereas the parameters for the two gA termini (formyl and ethanolamine) were the same as in [38] with the particular atom types renamed to conform to the CHARMM36 atom notation. For both the symmetric and asymmetric bilayers with gA, the system was first energy minimized for 10^4^ steps, then run for 100 ps with a 1 fs time step. Following this initial relaxation, the POPC/gA bilayer was simulated for 887 ns and the asymmetric bilayer with gA was simulated for 705 ns with a 2 fs time step. For more details on all simulated systems and the corresponding simulation parameters, see Section S1 in SM.

### Quantification of membrane deformation from simulations

To quantify the deformation of the membrane around gramicidin, the trajectories were first centered and aligned on the backbone of gA. Since gA can tilt with respect to the bilayer normal in the course of the simulations, special care was taken to ensure that the alignment step did not result in abnormal tilting of the membrane effectively leading to artificial changes in the bilayer thickness in the vicinity of the protein. The gA-lipid boundary was identified separately for each leaflet as the outermost layer of the time-averaged occupancy map (constructed on a 2 × 2 grid) of the respective gA monomer atoms projected onto the *xy* plane. The leaflet thickness at the gA-lipid boundary was calculated from the interpolated z-positions of the lipid atoms in the respective grid points, as described in Section S2 in SM. The leaflet thickness at the gA-lipid boundary was used to obtain the optimal deformation profile around a gA monomer by a self-consistent free energy minimization procedure as described in Section S3 in SM. The leaflet deformation around gA at distance *r* from the gA center was calculated as the squared deviation in thickness averaged over the area between the grid points at distance *r* from the gA center and the gA-lipid boundary (the bulk leaflet thickness was taken from the corresponding lipid-only simulations). The membrane deformation was taken as the sum of the deformations of the two leaflets.

### Quantification of membrane deformation in experiments

To estimate the amount of membrane deformation at each of the gA:lipid ratios used in the experiments, we first approximated the number *N* of gA dimers on a vesicle from: 1) the total surface area of a vesicle with diameter 130 nm (the average vesicle size measured with DLS); 2) the average area per lipid 64 Å^2^ (calculated from MD simulations for a POPC bilayer); and 3) the area per gA monomer 208 Å^2^ approximated from the occupancy map of the gA monomers described above. *N* gA dimers were then distributed uniformly on the surface of a sphere with diameter 130 nm using MATLAB, and the distance *d* between them was recorded. The membrane deformation at a given gA:lipid ratio was estimated as the membrane deformation at a distance *d*/2 from the gA center.

## RESULTS

### Gramicidin incorporation, conformation, and function are the same in symmetric and asymmetric liposomes

We modified a recently developed protocol for the construction of asymmetric lipid-only vesicles [30] to enable the preparation of asymmetric proteoliposomes. As described in Methods, the protocol involves mixing two populations of symmetric vesicles (only one of which contains pre-incorporated protein) and exchanging the lipids between their outer leaflets with MβCD (Fig. S1). Using this procedure we prepared two types of gramicidin-containing vesicles: 1) *compositionally symmetric* but *isotopically asymmetric* liposomes (s*LUVs) with an inner leaflet composed of POPC or its headgroup-perdeuterated variant (POPC-d13), and an outer leaflet exchanged with its chain-perdeuterated variant (POPC-d31); and 2) *compositionally asymmetric* vesicles (aLUV) with the same inner leaflet composition and an outer leaflet exchanged with the chain-perdeuterated variant of DMPC (DMPC-d54). Table S1 shows the donor mole fraction in the final samples as determined from GC/MS (see Methods) which ranged between 0.32 and 0.4 for the s*LUVs and between 0.35 and 0.45 for the aLUVs. The size of the vesicles was measured with DLS and was on average 130 nm in diameter with polydispersity index of < 0.2.

To examine the effect of the sample preparation protocol on the gramicidin, we assayed its incorporation, conformation, and function in the asymmetric proteoliposomes. Fig. 1A shows evidence of unaltered bilayer incorporation of gA from small-angle X-ray scattering (SAXS) data. In a SAXS experiment, hydrogen and deuterium scatter X-rays in the same way and the form factor of an isotopically asymmetric sample is indistinguishable from the form factor of the corresponding protiated sample. Moreover, because of the short length of gA relative to the bilayer thickness, larger amounts of the channel in the bilayer lead to systematic lift-off (i.e. increase in the intensity) of the first minimum in the SAXS form factor, as shown in the series of symmetric POPC LUVs with increasing gA mole fractions (Fig. 1A). Comparing (in Fig 1A) the isotopically asymmetric gA-containing LUVs (gA-s*LUV) prepared at a gA:lipid ratio of 1:40, with the series of symmetric LUVs, shows that the concentration of gA in the asymmetric LUVs is very close to the nominal mole fraction of gA initially incorporated in the acceptor vesicles; the incorporation was efficient and very little (if any) gA was lost during the asymmetric proteoliposome preparation.

**Figure 1.**
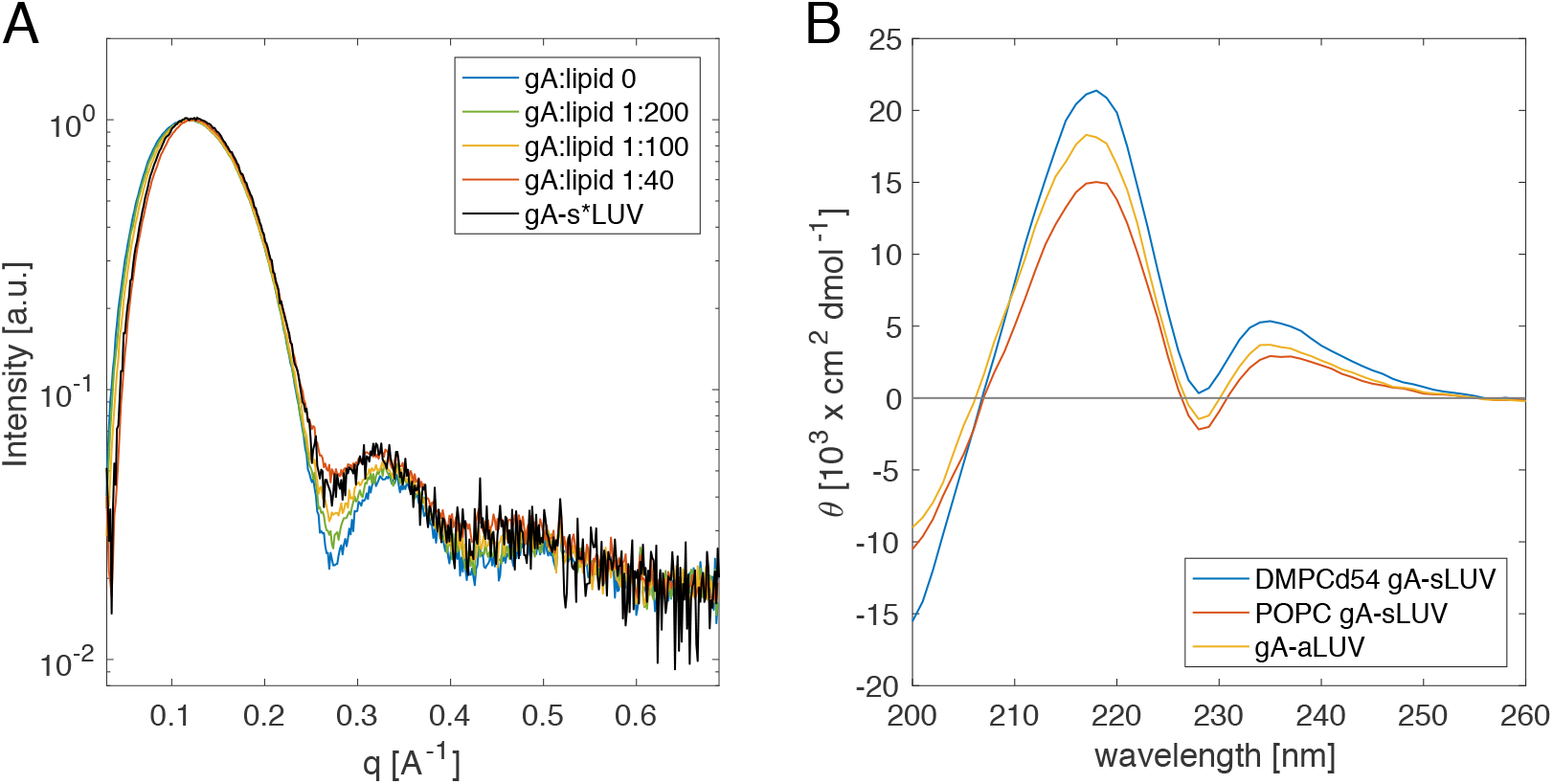
Gramicidin incorporation and conformation remain the same in symmetric and asymmetric liposomes. A) Color-coded SAXS form factors of a series of POPC gA-sLUVs with increasing concentration of gA and isotopically asymmetric LUVs, gA-s*LUV, composed of deuterated variants of POPC and prepared with a nominal gA:lipid ratio of 1:40. B) Circular dichroism spectra of gA-containing DMPC-d54 (blue) and POPC (red) sLUVs, and compositionally asymmetric LUVs composed of DMPC-d54 and POPC (yellow). All liposomes were prepared at a gA:lipid ratio of 1:40, and all measurements were performed at 25°C. See text for details.

Next we examined the structural properties of the incorporated gA. In addition to its canonical helical dimer conformation, gramicidin can exist in other non-functional conformations such as a dimeric helix in which two gA monomers are intertwined into a single extended helix [39]. The dimeric helix conformation can arise from hydrophobic mismatch (or from keeping gA in some nonpolar solvents) and has a distinct CD spectrum [40]. Fig 1B shows the CD spectrum of gA in the compositionally asymmetric LUVs in comparison with the spectra of gA in symmetric LUVs made of either just POPC or DMPCd54. The data show that most of the 0.025 mol% gA in the symmetric samples existed in a helical dimer conformation, and that the conformation remained unchanged during the steps of the gA-aLUVs preparation protocol.

To evaluate the function of gA in the asymmetric liposomes, we measured the rate with which a fluorophore (ANTS) trapped inside the vesicles was quenched by an externally added quencher (Thallium, Tl^+^). This assay provides a measure of the equilibrium dimer-to-monomer ratio of gA in the bilayer by taking advantage of the fact that Tl^+^ can enter the vesicles and quench ANTS only through functional gA dimers [29]. Fig. 2A shows the ANTS fluorescence decay as a function of time in the initially symmetric acceptor vesicles (66.1) and the final isotopically asymmetric gA-s*LUVs (62.9). According to the ratio of the two rates (0.95), the dimer-to-monomer equilibrium of gA in the isotopically asymmetric vesicles was essentially unaffected by the cyclodextrin-mediated lipid exchange. We obtained a similar ratio of 0.99 for the compositionally asymmetric liposomes (Fig. 2B), confirming that in both asymmetric samples gA function remained intact.

**Figure 2.**
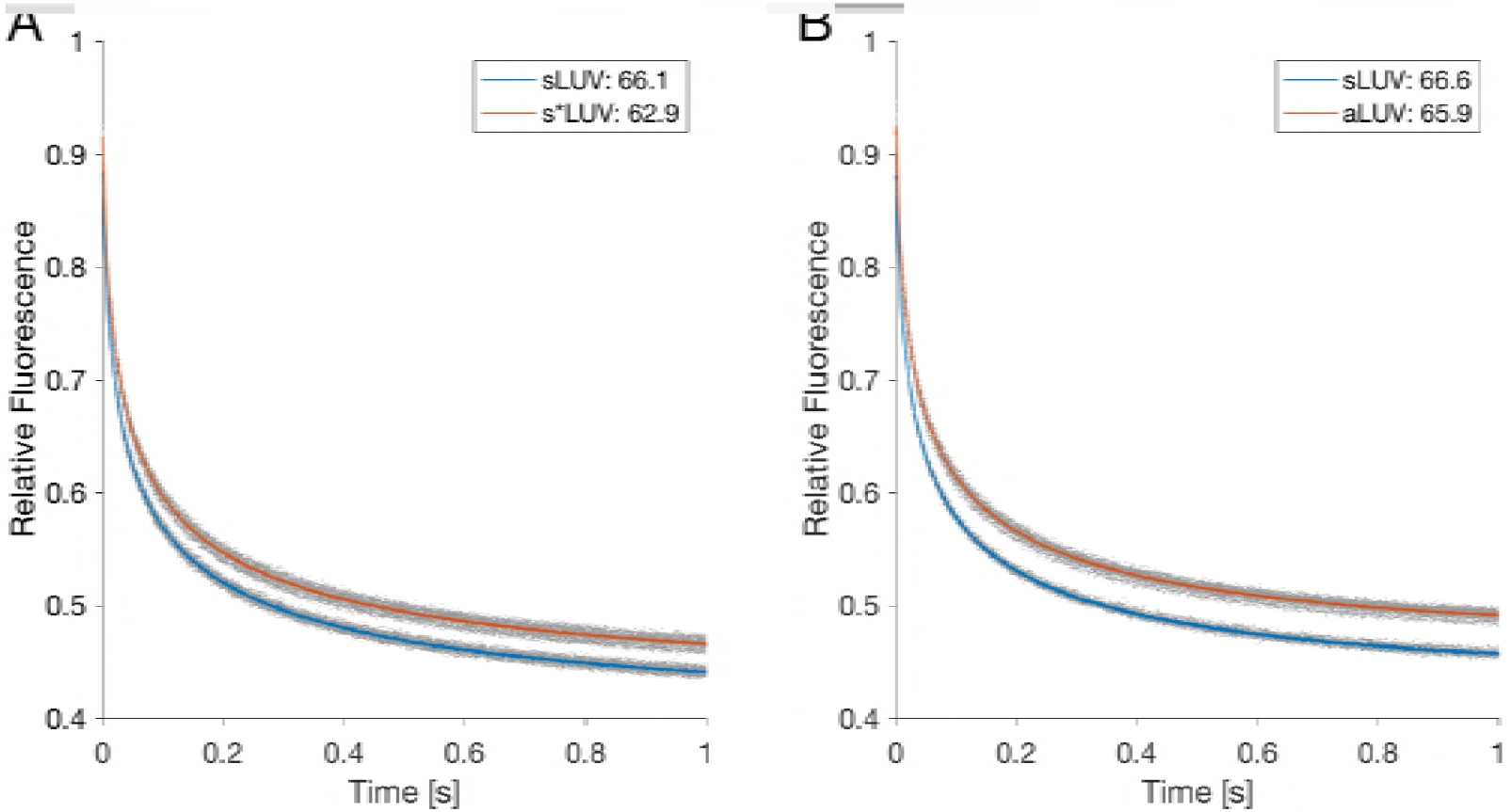
Gramicidin channel function remains intact in asymmetric liposomes. Changes in ANTS fluorescence over time for the isotopically (A) and compositionally (B) asymmetric samples (in red) and their corresponding symmetric acceptor vesicles (in blue). Replicate traces are shown in grey; their averages are shown in color. A constant background fluorescence measured before addition of Tl+ has been subtracted from all traces. The average rates calculated from the traces are indicated in the legends. All measurements were performed at ambient temperature of ~ 22°C.

### gA scrambles lipids in compositionally asymmetric vesicles

That gramicidin can scramble lipids first became evident in differential scanning calorimetry experiments. In DSC experiments, thermodynamic phase transitions (such as transitions between gel and fluid) are detected by monitoring the release of heat from a sample as a function of temperature. Fig. 3 shows a cooling temperature scan performed after sample preparation (scan 1) for compositionally asymmetric vesicles without (Fig. 3A) and with (Fig. 3B) gramicidin (see Methods). Two transition peaks are visible in both samples, likely coming from the different melting temperatures of the POPC-rich inner leaflet (POPC melting temperature is −2°C) and the DMPC-d54-rich outer leaflet (DMPC-d54 melting temperature is 19°C), see Fig. S3. After the first scan, the temperature was again brought to 30°C and a second cooling scan (scan 2) was performed without any observable changes in the DSC spectra, indicating that cycling through the temperature range of −8°C to 30°C (and consequently through the gel-liquid crystalline transition of each leaflet) by itself did not result in any major redistribution of the lipids between the two leaflets. In the gA-aLUV sample, however, after a set of fixed temperature incubations (20°C for 12 hrs, followed by 45°C for 5 h), the two peaks began to merge (scan 3, Fig. 3B) whereas those of the aLUV sample without gA remained unchanged (scan 3, Fig. 3A). The changes in the gA-aLUV sample were even more pronounced after subjecting the samples to another set of fixed temperature incubations (scan 4). These results indicate that gramicidin facilitated the exchange of lipid between the two leaflets, presumably through its ability to accelerate lipid flip-flop, which eventually would result in a symmetric bilayer with a single transition temperature peak (grey dashed lines in Fig. 3).

**Figure 3.**
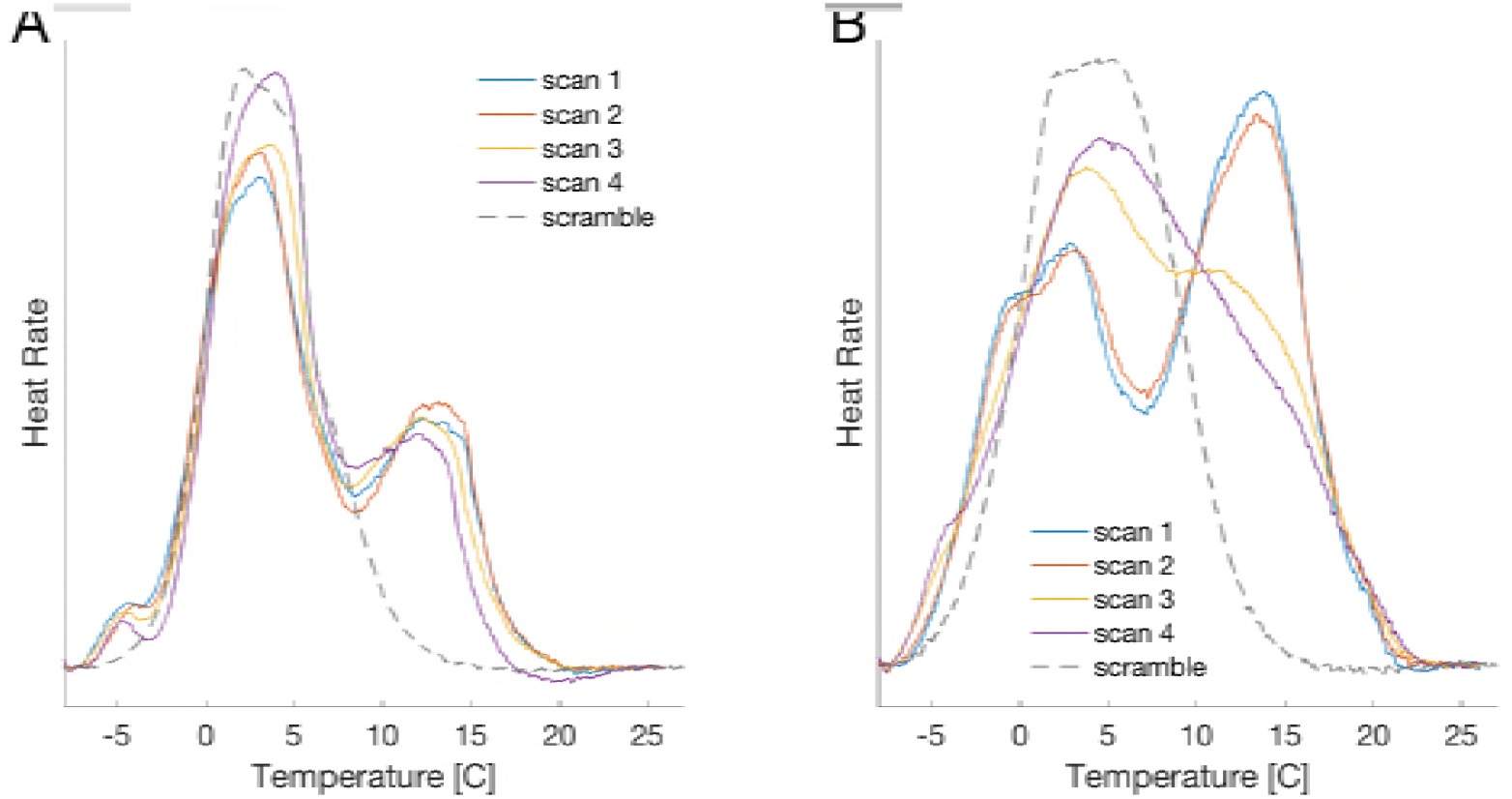
Gramicidin scrambles lipids in compositionally asymmetric vesicles. DSC data of compositionally asymmetric LUVs with DMPC-d54 and POPC, and without (A) or with (B) gramicidin at gA:lipid ratio of 1:40. Four consecutive cooling scans were performed as follows: after the asymmetric vesicle preparation (scan 1, blue); after scan 1 (scan 2, red); after subsequent incubation at 20°C for 12 h followed by incubation at 45°C for 5 h (scan 3, yellow); and after another set of incubations at 20°C for 12 h and 45°C for 5 h (scan 4, purple). Also shown for comparison with grey dashed lines is data for the symmetric samples (scramble) with the same overall composition as the asymmetric vesicles (Table S1).

### gA increases the rate of lipid flip-flop in compositionally asymmetric vesicles

To quantify the effect of gA on lipid flip-flop, we used ^1^H NMR to measure the rate of lipid translocation in the presence and absence of the channel (see Methods). The nine equivalent protons on a protiated choline headgroup produce a clearly defined resonance peak at 3.1-3.6 ppm. The shift reagent Pr^3+^, when added externally to the sample, interacts only with the lipid headgroups exposed on the outer leaflet of the vesicles and shifts their resonance downfield [41, 42]. After exchanging lipids with protiated choline headgroups (POPC-d31 or DMPC-d54) into acceptor vesicles composed of the headgroup-deuterated POPC-d13, the only detectable resonance comes from the exchanged donor lipids. The distribution of the protiated-headgroup lipids across the two bilayer leaflets thus can be determined from the ratio of the areas of the shifted (*C_out_*) and unshifted (*C_in_*) choline peaks in the NMR spectrum. Repeating the shift experiment on different aliquots of the sample over time allows for direct quantification of the lipid flip-flop rate [42]. The sample was thus incubated at room temperature in the absence of Pr^3+^; the shift reagent was used only during the measurement as described in Materials and Methods.

Fig. 4 shows the relative changes in transverse lipid distribution Δ*C* (see Eq. 2) as a function of time *t* for both the chemically symmetric (Fig. 4A) and asymmetric (Fig. 4B) liposomes with different amounts of gramicidin (see Methods). Table 2 lists the corresponding lipid flip-flop rates and half-times calculated as described in Methods. Whereas the spontaneous lipid flip-flop rate in the absence of gA is immeasurably slow in the s*LUVs, it is visibly increased when DMPC is exchanged into the outer leaflet of POPC. Importantly, the rate of lipid translocation in both samples increases in the presence of the channel, in a gA-concentration dependent manner.

**Figure 4.**
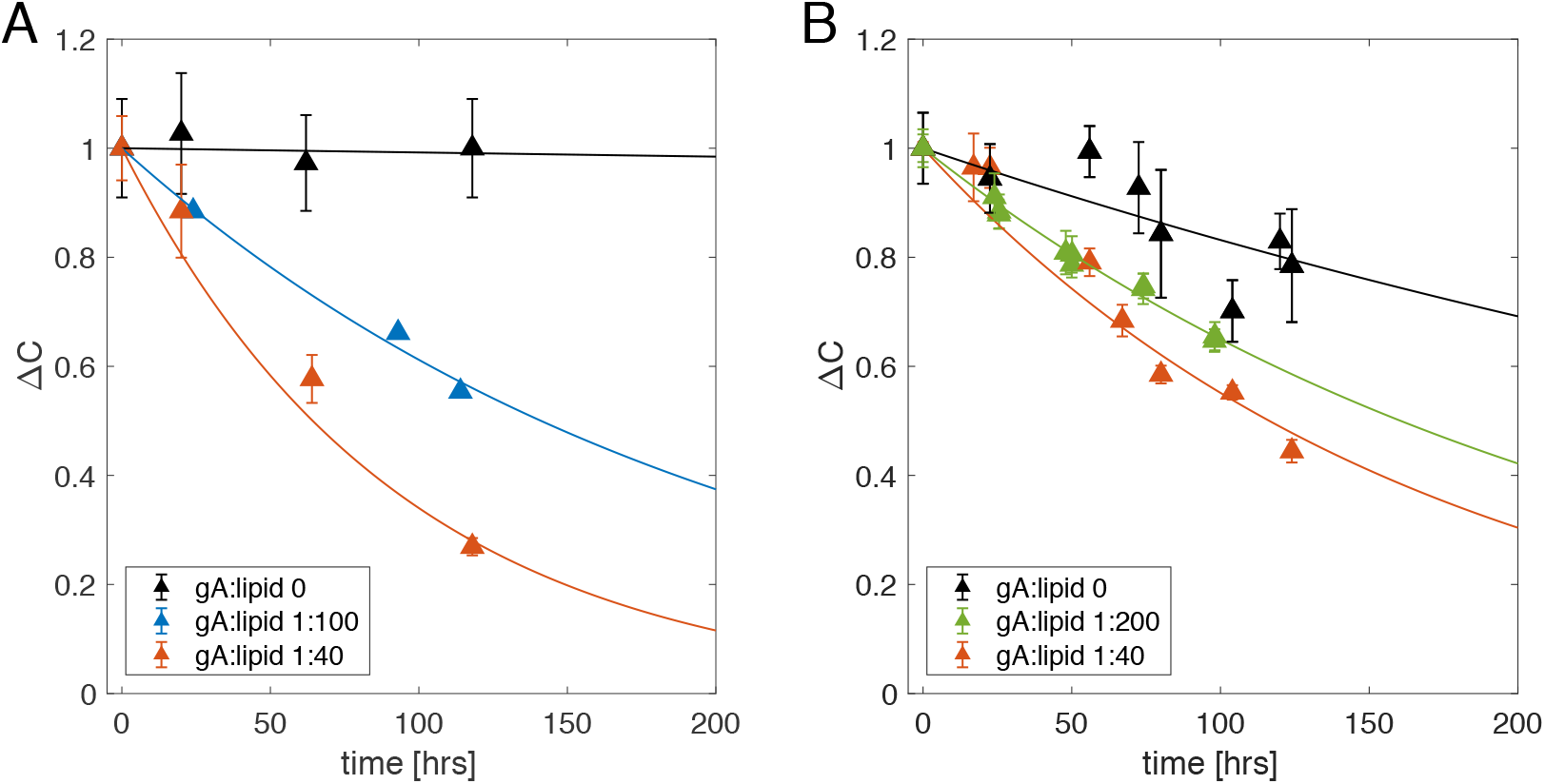
Gramicidin increases the rate of lipid flip-flop in isotopically and compositionally asymmetric vesicles. Time evolution of the interleaflet distribution of POPC-d31 in s*LUVs (A) and DMPC-d54 in aLUVs (B) either without gA (black) or with gA at gA:lipid ratio of 1:40 (red), 1:100 (blue) and 1:200 (green). Both plots show the time-dependent changes in the distribution of POPC-31 between the outer and inner leaflets, relative to the first time point measured after vesicle preparation (ΔC). See text for details. Error bars represent standard deviations of at least 3 consecutive Pr^3+^ additions. Each time trace in panel A is from a single sample. The time traces of the compositionally asymmetric vesicles in panel B are from 1 (gA:lipid 1:40), 2 (no gA) and 3 (gA:lipid 1:200) separately prepared samples, respectively. The kinetics reported here correspond to sample behavior at an ambient temperature of ~ 22°C.

**Table 2.** Translocation kinetics in the examined systems. Shown are the corresponding half times and rates of lipid flip-flop calculated from the NMR data shown in Fig. 4 as described in Methods. The numbers in brackets represent 95% confidence intervals.

| Outer leaflet | gA:lipid | $t_{1/2}$ [h] | $k_f / 10^{-7} s^{-1}$ |
| --- | --- | --- | --- |
| POPC-d31 / POPC-d13 | 0 | ~ 8900 | ~ 0.1 |
|  | 1:100 | 141 [125; 163] | 6.8 [5.9; 7.7] |
|  | 1:40 | 64 [55; 76] | 16.0 [12.6; 17.4] |
| DMPC-d54 / POPC-d13 | 0 | 376 [251; 748] | 2.6 [1.3; 3.8] |
|  | 1:200 | 160 [153; 168] | 6.0 [5.7; 6.3] |
|  | 1:40 | 116 [104; 132] | 8.3 [7.3; 9.2] |

### Models for gA-mediated lipid flip-flop

If every gA dimer accelerated the movement of lipid between the leaflets, the rate of lipid flip-flop would vary linearly with gA mole fraction [20]. Fig. 5 shows the flip-flop rates calculated from the NMR data (Table 2), as a function of the gA:lipid ratio. Indeed, the data for the s*LUV samples (shown in blue) are consistent with the model of single gA channel-mediated lipid translocation. In the aLUV samples (shown in red), however, the linear relationship predicted from the model is not as clear, and the uncertainty in the data precludes any firm conclusions. Considered together, the two data sets introduce a conundrum: in the absence of gramicidin, the flip-flop rate in the compositionally asymmetric bilayers is clearly faster than the rate in the symmetric membranes, yet in the presence of gA at gA:lipid ratio of 1:40 this trend is reversed and the flip-flop kinetics in the aLUVs are much slower than those in the s*LUVs. This result suggests a mechanism different from a single channel-induced perturbation by which gramicidin catalyzes lipid translocation, based on the following considerations: 1) as the gA channel is relatively short, it is likely to cause thickness deformations in the membrane; and 2) the thickness deformations in a POPC bilayer could be alleviated in the presence of a shorter-chain lipid like DMPC. Thus gA-induced bilayer stress could be a plausible contributing factor to the observed trends.

**Figure 5.**
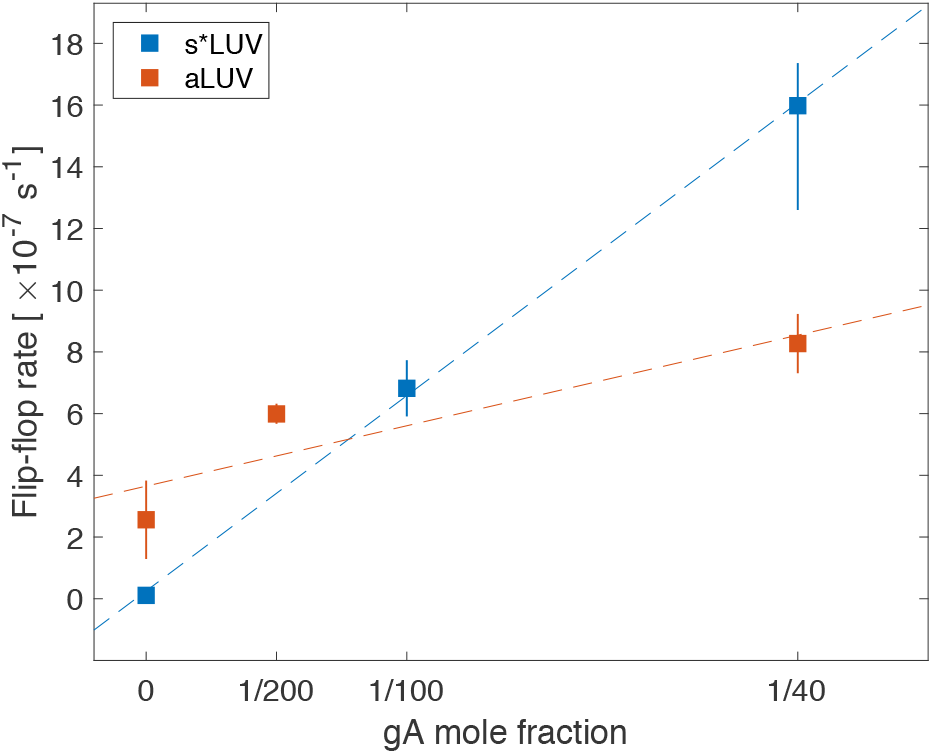
Rate of lipid flip-flop shows a complex relationship with gA mole fraction. Flip-flop rates calculated from the data in Fig. 4 for the compositionally symmetric (blue) and asymmetric (red) LUVs (see Table 2) as a function of the nominal gA mole fraction in the samples. Error bars represent 95% confidence intervals.

### Membrane deformations are likely to play a role in gA-mediated lipid flip-flop

Gramicidin has been routinely used as a model peptide to study local bilayer deformations that result from protein-membrane interactions [43, 44]. It is in this context important that the gA channel’s helical pitch is invariant with respect to changes in the channel-bilayer hydrophobic mismatch, meaning that bilayer will adjust to the channel rather than the channel adjusting to the bilayer (in bilayers having a hydrophobic thickness greater than the channel length) [45]. gA has been shown to induce thinning in a pure DMPC bilayer [46] and one would expect that the same effect would be observed in bilayers with hydrophobic thickness greater than DMPC, such as a membrane composed of POPC [47]. Such local deformations produce a stress on the membrane that will increase with increasing gA concentration in the membrane and could potentially affect lipid flip-flop. We therefore investigated the response of the bilayer to the presence of a single gA channel in the two experimentally studied systems.

MD simulations, in combination with the continuum theory of membrane deformations (CTMD) [48], were used to estimate the amount of membrane deformation at the different gA mole fractions from the experiments by first quantifying the changes in membrane thickness as a function of distance from the channel, and then relating them to the average distances between channels in the samples. To enable this protocol, MD simulations were performed on a system containing a single gA channel embedded in a symmetric POPC bilayer, or in an asymmetric membrane with a top leaflet of DMPC/POPC 0.75/0.25 and a bottom leaflet of DMPC/POPC 0.10/0.90. The compositions of the two leaflets were based on a different set of samples prepared for small-angle neutron scattering experiments (unpublished), and were similar to the leaflet compositions of the samples examined with NMR in the present work (Table S1). At the end of the simulations, the thickness of each leaflet at the gA-lipid boundary was analyzed and used to calculate the optimal thickness deformation profile around gA using the CTMD formalism, as described in Methods and Section S3 in SM. Because the membrane deformation energy varies with the square of the hydrophobic mismatch (see Eq. S3 in SM), Fig. 6A shows the resulting average squared deviations in thickness as a function of distance from the gA center in both systems. The amount of membrane deformation decreases gradually further away from the protein in the two bilayers, and it is less in the asymmetric membrane, indicating that gA is able to adapt more easily to the asymmetric bilayer environment.

**Figure 6.**
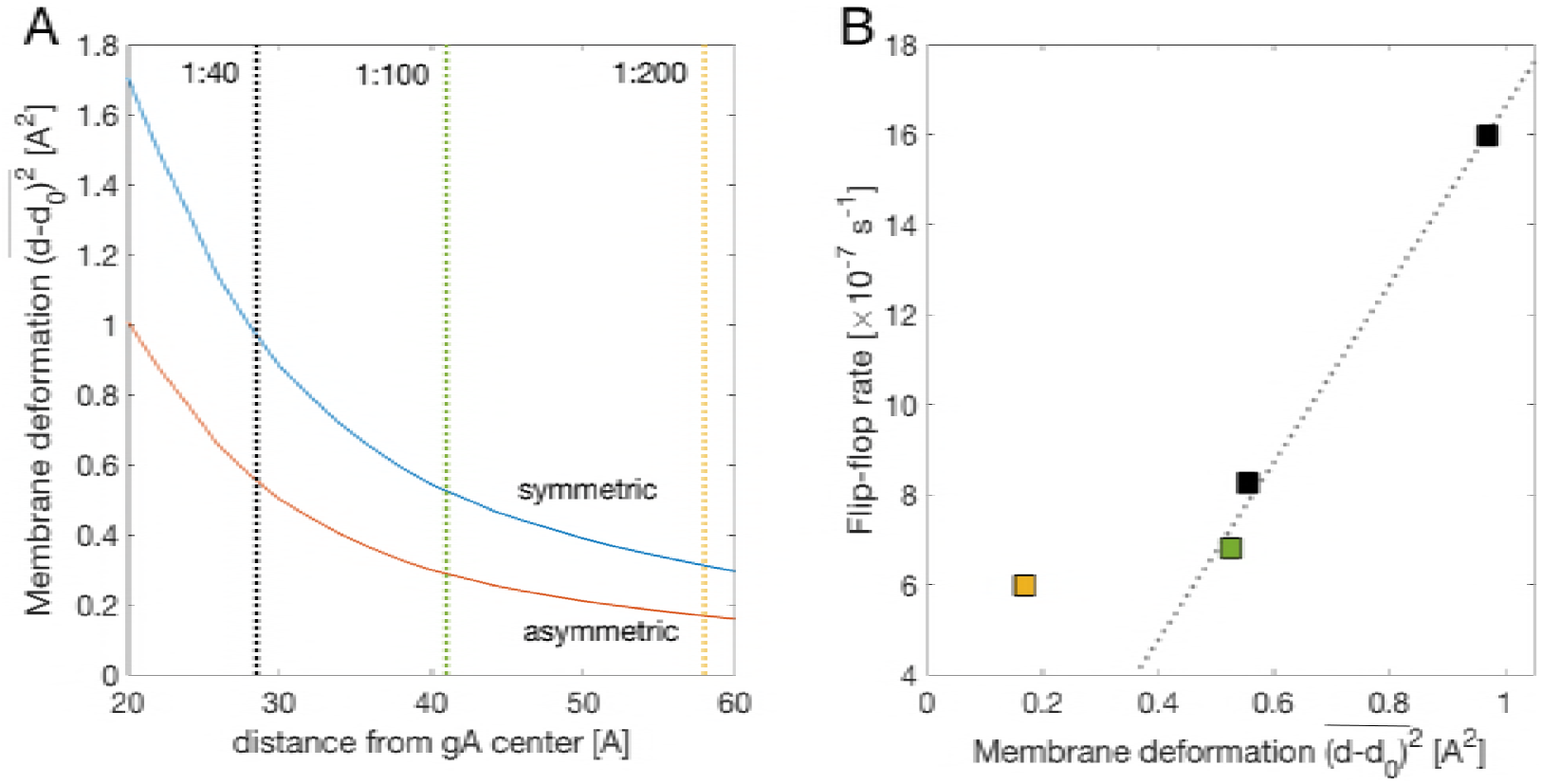
The gA-mediated lipid flip-flop rate at high gA concentrations correlates with the membrane deformation. (A) Average squared deviations in bilayer thickness as a function of distance from gA center for a symmetric POPC bilayer (blue) and an asymmetric bilayer like in the NMR experiments in Fig. 4 (red). The thickness deviations were calculated from the membrane deformation profiles around a single gA channel obtained by a CTMD-guided free energy minimization utilizing information from MD simulations (see Methods). Dashed lines indicate the approximated half-distances between gA channels on the surface of the LUVs at three different gA:lipid ratios: 1:40 (black), 1:100 (green) and 1:200 (yellow). (B) gA-mediated lipid flip-flop rate from Fig. 5 as a function of the corresponding membrane deformation from panel (A). The data points are color coded according to gA concentration as in panel (A).

To examine whether membrane deformations could be involved in the ability of gramicidin to scramble lipids, we approximated the packing density of gA dimers within the experimentally prepared LUVs. We estimated the distance between the channels at each gA mole fraction by assuming that they are uniformly distributed on the surface of the vesicles (see Methods). The dashed lines on Fig. 6A indicate the respective half distances between the gA centers. The compositionally symmetric sample with gA at gA:lipid ratio of 1:40 in which the rate of lipid flip-flop was highest (Table 2) also seemed to have the largest membrane deformation among the examined asymmetric proteoliposomes. Fig. 6B shows the relationship between the measured rates of lipid flip-flop in the four gA-aLUVs and the corresponding amounts of membrane deformation as quantified by the analysis in Fig. 6A. Whereas the correlation between gA-mediated lipid scrambling and membrane deformation at gA mole fractions of 0.01 and higher is very strong (0.998), the sample with gA:lipid ratio of 1:200 did not follow the same relationship and appeared as an outlier (Fig. 6B). This result suggests that the stress induced in the membrane as a result of gA-membrane interactions is an important contributor to the observed effects of gA on lipid scrambling, and that at high mole fractions the channels accelerate lipid flip-flop in a mechanistically different way than at low gA concentrations.

## DISCUSSION

In the study of protein-membrane interactions, the effects of the membrane (e.g. its composition and structure) on protein function and oligomerization have been examined extensively. It is equally important, however, to account for the protein’s effect on the membrane, because a protein embedded in a bilayer is prone to introduce defects that can perturb the bilayer structure and affect its transverse and lateral organization. We have, therefore, developed new protocols for the systematic investigation of protein-mediated changes in the lipid compositions of the two bilayer leaflets by utilizing model systems that allow for fine tuning of various membrane parameters. As discussed below, these protocols have been devised to exploit specific biophysical properties that bring to light specific elements of the complex interplay between the protein and the membrane, and to quantify the consequences.

We first discuss the preparation and properties of the gA-containing asymmetric lipid vesicles; we then discuss the effect of gA on lipid flip-flop; and, finally, we present a possible mechanism for the gA-induced lipid scrambling.

### Preparation and biophysical characterization of asymmetric proteoliposomes

Asymmetric membranes containing transmembrane (TM) proteins have been successfully prepared in the past. Two general types of approaches have been adopted. In one, the protein (either soluble or micelle-stabilized) is added externally to pre-formed asymmetric membranes including LUVs filled with sucrose [49], droplet interface bilayers [50] and planar supported bilayers [51]. In the other, also employed here, the protein is first reconstituted into symmetric vesicles, then the outer leaflet of the vesicles is exchanged for specific lipids. For example, following the latter approach Vitrac et al. prepared sonicated small unilamellar vesicles (SUVs) with the 12-transmembrane protein LacY from *E. coli* and examined the effect of asymmetric charge distribution on the topology of the protein [4, 52]. In a different study, asymmetric SUVs with the nicotinic acetylcholine receptor (AChR) were prepared by using MβCD-loaded lipid complexes to exchange the lipids on the outer leaflet of symmetric SUVs containing AChR, with sphingomyelin [3]. While these examples illustrate how a variety of TM proteins can be incorporated into asymmetric model membranes of different geometries, the effect of the protein (and protein incorporation) on the lipid compositions of the two bilayer leaflets can vary and therefore has to be examined explicitly. With respect to this effect, the protocols and assays presented here constitute an advantageous platform for the biophysical characterization of the sample, much improved by utilizing large tensionless proteoliposomes free from either the potential effects of curvature, or the commonly found residues of organic solvents, or polymer cushion supports. Thus, we found only minimal effects of the preparation protocol on the incorporation, conformation and function of gramicidin (Figs. 1 and 2), but these are likely to depend on the types of lipids in the vesicles, and their negligible magnitude cannot be taken for granted. Indeed, the energy cost of gA dimer formation increases with increasing bilayer thickness, meaning that the monomer-to-dimer equilibrium of gA is shifted towards the monomeric state [40, 53], and in such cases the gA monomers have been shown to more easily exchange between vesicles [54]. This could result in a loss of the protein during the CD-mediated lipid exchange between donor and acceptor vesicles. Furthermore, a preference of a gA monomer for the composition of one leaflet vs. another in the gA-aLUVs may require additional tests of the interleaflet gA localization and vesicle stability. The thermodynamic phase properties of the bilayer in the presence of gA (whether in dimeric or monomeric state), should also be considered. For example, we found that gA broadens and slightly lowers the transition temperature of DMPCd54 (Fig. S4), but this effect is likely to depend on both the protein type and concentration in the membrane, and may be different for other proteins. A gel environment could affect not only the protein dynamics but also the efficiency of lipid exchange during the aLUV preparation protocol (see [28]).

### Rates of lipid flip-flop measured with NMR

The slow rate of spontaneous lipid flip-flop that we measured in the chemically symmetric vesicles in the absence of gA is consistent with previous reports [14, 55]. The flip-flop kinetics in the compositionally asymmetric membranes, however, were significantly faster (Fig. 5). Because both POPC and DMPC have phosphocholine headgroups, this increase is likely due to differences in the chain lengths of the lipids (16 and 18 carbons for POPC and 14 carbons for DMPC) and the corresponding leaflet thicknesses (Table S2). This explanation is consistent with results from small-angle neutron scattering experiments showing a faster flip-flop rate in DMPC LUVs compared to POPC LUVs [55], and the corresponding free energy for flip-flop in the two bilayers, quantified with MD simulations [56].

Our results for the gA-mediated half time of lipid translocation in the POPC liposomes (t_1/2_ ≈ 64 h at a gA:lipid ratio of 1:40) are comparable to those of Fattal and coworkers with the chain-labeled fluorescent lipid analog NBD-PC (*t*_1/2_ > 96 h) [14]. This similarity is surprising in light of the different chemical structures of the acyl chains of NBD-PC relative to POPC and suggests that, in agreement with earlier experimental observations [57], the nature of the lipid headgroup is the dominant factor in determining the flip-flop rate. Yet, the chain structure also has been shown to strongly affect flip-flop for lipids with multiple double bonds [58]. In contrast, the reported kinetics of lipid translocation in the presence of gA in erythrocytes [24] and supported lipid bilayers [25] are much faster, likely due to the specific experimental conditions in the studies as discussed in the Introduction.

The lipid flip-flop rates accessible to NMR measurements are limited by the time needed to perform a single shift experiment. Depending on the type of instrument (i.e. the strength of the magnet) and sample concentration, one measurement in the absence of shift reagent, followed by 3 Pr^3+^ titrations (needed to obtain error bars, for example) could take anywhere from 15 min to 1 hour. Thus this technique can be used to measure only slower time-dependent translocation processes. However, since the lipid flip-flop rate would be expected to increase with increasing temperatures, systems with expected faster rates can be compared if incubated at lower temperatures, see also [59]. The samples in this study were incubated on the benchtop at an ambient temperature of ~22°C.

### Mechanisms of gA-induced lipid scrambling

It has previously been proposed that the ability of gA to scramble lipid analogs is not due to bilayer perturbations induced by individual gA channels because [23, 24]: 1) gA increases the translocation rate of lysophosphatidylcholine only at concentrations much higher than those where gA performed its normal function as a conducting channel, but where gA produces non-specific solute leakage; and 2) formylation of gA’s tryptophan residues, which prevents formation of gA clusters, abolished the gA-catalyzed lyso-lipid scrambling. Instead, it was proposed that gA at gA:lipid mole fractions of 1:2000 or higher forms some sort of aggregate(s) in the erythrocyte membrane, which may be intermediates for the formation of hexagonal phases. The proposed clusters would induce transient defects in the bilayer with subsequent formation of aqueous leaks that allow for the passage of molecules as large as choline and sucrose across the cell membrane, as well as the translocation of lipid analogs [24]. The proposed formation of gA clusters would depend on channel-bilayer hydrophobic mismatch because gA did not cause detectable aggregates in DMPC bilayers, where there is minimal channel-bilayer hydrophobic mismatch [46] and little accumulation of stress in the membrane. Though we cannot exclude the existence of gA aggregates in our samples, their presence is unlikely [46, 60] and the lamellar SAXS form factors (Fig. 1A) definitively exclude the presence of non-lamellar phases known to form at high gA concentrations under different experimental conditions [19]. Our NMR and computational analysis further illuminate the importance of the frustration energy in the bilayer: At high channel densities, the bilayer thickness is not able to relax to its unperturbed state between the channels (Fig. 6), leading to bilayer-deformation-induced stress. This stress would increase the probability of transient clusters of bilayer-spanning gA channels, e.g. [61, 62], which could serve to decrease the energy barrier for lipid translocation and thereby increase the lipid flip-flop rate [63]. The deviation of the gA-aLUVs at gA:lipid ratio of 1:200 from the other samples in Fig. 6B further suggests that at this lower gA mole fraction the role of the frustration energy is different, resulting in a mechanistically distinct route for the observed gA effect.

## CONCLUSIONS

Using new methodologies and protocols for preparing and characterizing asymmetric proteoliposomes to the system of gramicidin channels, we have demonstrated the ability of gA to accelerate inter-leaflet lipid translocation in both chemically symmetric and asymmetric membranes. The mechanistic analysis of the results show that the channel-induced bilayer deformation likely contributes to the rate of lipid flip-flop. The ability to characterize and quantify the interplay between transmembrane proteins and their solvating lipid environment allows us to determine the properties of the protein-laden bilayers. If such properties are considered properly, e.g., with the type of methodology illustrated here, the mechanistic understanding of much more complex biomimetic systems becomes feasible and practical.

## Author Contributions

MD, FAH, HW, GWF and OSA designed the research. MD, FAH, RR and LS performed the experiments. MD, FAH, DM and TP analyzed the experimental data. MD performed all simulations and computational analysis. MD, FAH, DM, JK, GWF, HW and OSA wrote the manuscript.

## ACKNOWLEDGEMENTS

We thank Haden Scott for technical assistance with CD measurements, Andrew Beaven for providing the MD force field parameters for gramicidin-A, and W. Clay Bracken for help with identifying the water suppression conditions for the NMR measurements at Weill Cornell Medical College. This work was supported by NSF grant No. MCB-1817929 (FAH), NIH grants P01 DA012408 (HW) and R01 GM021342 (OSA), and by a grant from Natural Sciences and Engineering Research Council of Canada (NSERC), [funding reference number 2018-04841] (DM). GC/MS, DSC, and DLS measurements were supported by the Biophysical Characterization Laboratory suite of the Shull Wollan Center at Oak Ridge National Laboratory (ORNL). SAXS measurements were supported by DOE scientific user facilities at ORNL.

## Reference

1. Collins, M.D. and S.L. Keller, Tuning lipid mixtures to induce or suppress domain formation across leaflets of unsupported asymmetric bilayers. Proc Natl Acad Sci U S A, 2008. 105(1): p. 124–8.

2. Kiessling, V., C. Wan, and L.K. Tamm, Domain coupling in asymmetric lipid bilayers. Biochim Biophys Acta, 2009. 1788(1): p. 64–71.

3. Perillo, V.L., et al., Transbilayer asymmetry and sphingomyelin composition modulate the preferential membrane partitioning of the nicotinic acetylcholine receptor in Lo domains. Arch Biochem Biophys, 2016. 591: p. 76–86.

4. Vitrac, H., et al., Dynamic membrane protein topological switching upon changes in phospholipid environment. Proc Natl Acad Sci U S A, 2015. 112(45): p. 13874–9.

5. Perlmutter, J.D. and J.N. Sachs, Interleaflet interaction and asymmetry in phase separated lipid bilayers: molecular dynamics simulations. J Am Chem Soc, 2011. 133(17): p. 6563–77.

6. St. Clair, J.R., et al., Preparation and Physical Properties of Asymmetric Model Membrane Vesicles, in The Biophysics of Cell Membranes. Springer Series in Biophysics, R. Epand and J.M. Ruysschaert, Editors. 2017, Springer: Singapore.

7. Markones, M., et al., Engineering Asymmetric Lipid Vesicles: Accurate and Convenient Control of the Outer Leaflet Lipid Composition. Langmuir, 2018. 34(5): p. 1999–2005.

8. Takaoka, R., et al., Formation of asymmetric vesicles via phospholipase D-mediated transphosphatidylation. Biochim Biophys Acta, 2018. 1860(2): p. 245–249.

9. Contreras, F.X., et al., Transbilayer (flip-flop) lipid motion and lipid scrambling in membranes. FEBS Lett, 2010. 584(9): p. 1779–86.

10. Sperotto, M.M. and A. Ferrarini, Spontaneous Lipid Flip-Flop in Membranes: A Still Unsettled Picture from Experiments and Simulations, in The Biophysics of Cell Membranes. Springer Series in Biophysics, R. Epand and J.M. Ruysschaert, Editors. 2017, Springer: Singapore.

11. Pomorski, T.G. and A.K. Menon, Lipid somersaults: Uncovering the mechanisms of protein-mediated lipid flipping. Prog Lipid Res, 2016. 64: p. 69–84.

12. Sebastian, T.T., et al., Phospholipid flippases: building asymmetric membranes and transport vesicles. Biochim Biophys Acta, 2012. 1821(8): p. 1068–77.

13. Aye, I.L., A.T. Singh, and J.A. Keelan, Transport of lipids by ABC proteins: interactions and implications for cellular toxicity, viability and function. Chem Biol Interact, 2009. 180(3): p. 327–39.

14. Fattal, E., et al., Pore-forming peptides induce rapid phospholipid flip-flop in membranes. Biochemistry, 1994. 33(21): p. 6721–31.

15. Tieleman, D.P. and S.J. Marrink, Lipids out of equilibrium: energetics of desorption and pore mediated flip-flop. J Am Chem Soc, 2006. 128(38): p. 12462–7.

16. Ernst, O.P. and A.K. Menon, Phospholipid scrambling by rhodopsin. Photochem Photobiol Sci, 2015. 14(11): p. 1922–31.

17. Andersen, O.S. and R.E. Koeppe, 2nd, Bilayer thickness and membrane protein function: an energetic perspective. Annu Rev Biophys Biomol Struct, 2007. 36: p. 107–30.

18. de Kruijff, B., E.J. van Zoelen, and L.L. van Deenen, Glycophorin facilitates the transbilayer movement of phosphatidylcholine in vesicles. Biochim Biophys Acta, 1978. 509(3): p. 537–42.

19. Tournois, H., et al., Gramicidin-induced hexagonal HII phase formation in erythrocyte membranes. Biochemistry, 1987. 26(21): p. 6613–21.

20. Kol, M.A., et al., Membrane-spanning peptides induce phospholipid flop: a model for phospholipid translocation across the inner membrane of E. coli. Biochemistry, 2001. 40(35): p. 10500–6.

21. Kol, M.A., et al., Phospholipid flop induced by transmembrane peptides in model membranes is modulated by lipid composition. Biochemistry, 2003. 42(1): p. 231–7.

22. Anglin, T.C., K.L. Brown, and J.C. Conboy, Phospholipid flip-flop modulated by transmembrane peptides WALP and melittin. J Struct Biol, 2009. 168(1): p. 37–52.

23. Killian, J.A., Gramicidin and gramicidin-lipid interactions. Biochim Biophys Acta, 1992. 1113(3-4): p. 391–425.

24. Classen, J., et al., Gramicidin-induced enhancement of transbilayer reorientation of lipids in the erythrocyte membrane. Biochemistry, 1987. 26(21): p. 6604–12.

25. Anglin, T.C., J. Liu, and J.C. Conboy, Facile lipid flip-flop in a phospholipid bilayer induced by gramicidin A measured by sum-frequency vibrational spectroscopy. Biophys J, 2007. 92(1): p. L01–3.

26. Marquardt, D., et al., (1)H NMR Shows Slow Phospholipid Flip-Flop in Gel and Fluid Bilayers. Langmuir, 2017. 33(15): p. 3731–3741.

27. Kucerka, N., et al., Curvature effect on the structure of phospholipid bilayers. Langmuir, 2007. 23(3): p. 1292–9.

28. Doktorova, M., et al., Preparation of asymmetric phospholipid vesicles for use as cell membrane models. Nature Protocols, 2018. In press. DOI: 10.1038/s41596-018-0033-6

29. Ingolfsson, H.I. and O.S. Andersen, Screening for small molecules’ bilayer-modifying potential using a gramicidin-based fluorescence assay. Assay Drug Dev Technol, 2010. 8(4): p. 427–36.

30. Heberle, F.A., et al., Subnanometer Structure of an Asymmetric Model Membrane: Interleaflet Coupling Influences Domain Properties. Langmuir, 2016. 32(20): p. 5195-200.

31. Jo, S., et al., CHARMM-GUI: a web-based graphical user interface for CHARMM. J Comput Chem, 2008. 29(11): p. 1859–65.

32. Jo, S., et al., CHARMM-GUI Membrane Builder for mixed bilayers and its application to yeast membranes. Biophys J, 2009. 97(1): p. 50–8.

33. Lee, J., et al., CHARMM-GUI Input Generator for NAMD, GROMACS, AMBER, OpenMM, and CHARMM/OpenMM Simulations Using the CHARMM36 Additive Force Field. J Chem Theory Comput, 2016. 12(1): p. 405–13.

34. Doktorova, M. and H. Weinstein, Accurate in silico modeling of asymmetric bilayers based on biophysical principles. Submitted.

35. Phillips, J.C., et al., Scalable molecular dynamics with NAMD. J Comput Chem, 2005. 26(16): p. 1781–802.

36. Klauda, J.B., et al., Improving the CHARMM force field for polyunsaturated fatty acid chains. J Phys Chem B, 2012. 116(31): p. 9424–31.

37. Klauda, J.B., et al., Update of the CHARMM all-atom additive force field for lipids: validation on six lipid types. J Phys Chem B, 2010. 114(23): p. 7830–43.

38. Beaven, A.H., et al., Gramicidin A Channel Formation Induces Local Lipid Redistribution I: Experiment and Simulation. Biophys J, 2017. 112(6): p. 1185–1197.

39. Bystrov, V.F. and A.S. Arsenev, Diversity of the gramicidin A spatial structure: Twodimensional 1HNMR study in solution. Tetrahedron, 1988. 44(3): p. 925–940.

40. Galbraith, T.P. and B.A. Wallace, Phospholipid chain length alters the equilibrium between pore and channel forms of gramicidin. Faraday Discuss, 1998(111): p. 159–64; discussion 225–46.

41. Perly, B., et al., Estimation of the location of natural alpha-tocopherol in lipid bilayers by 13C-NMR spectroscopy. Biochim Biophys Acta, 1985. 819(1): p. 131–5.

42. Marquardt, D., et al., 1H NMR Shows Slow Phospholipid Flip-Flop in Gel and Fluid Bilayers. Langmuir, 2017. 33(15): p. 3731–3741.

43. Andersen, O.S., et al., Single-molecule methods for monitoring changes in bilayer elastic properties. Methods Mol Biol, 2007. 400: p. 543–70.

44. Lundbaek, J.A., et al., Lipid bilayer regulation of membrane protein function: gramicidin channels as molecular force probes. J R Soc Interface, 2010. 7(44): p. 373–95.

45. Katsaras, J., et al., Constant helical pitch of the gramicidin channel in phospholipid bilayers. Biophysical journal, 1992. 61(3): p. 827–830.

46. Harroun, T.A., et al., Experimental evidence for hydrophobic matching and membrane-mediated interactions in lipid bilayers containing gramicidin. Biophys J, 1999. 76(2): p. 937–45.

47. Kucerka, N., M.P. Nieh, and J. Katsaras, Fluid phase lipid areas and bilayer thicknesses of commonly used phosphatidylcholines as a function of temperature. Biochim Biophys Acta, 2011. 1808(11): p. 2761–71.

48. Mondal, S., et al., Quantitative modeling of membrane deformations by multihelical membrane proteins: application to G-protein coupled receptors. Biophys J, 2011. 101(9): p. 2092–101.

49. Lin, Q. and E. London, The influence of natural lipid asymmetry upon the conformation of a membrane-inserted protein (perfringolysin O). J Biol Chem, 2014. 289(9): p. 5467-78.

50. Hwang, W.L., et al., Asymmetric droplet interface bilayers. J Am Chem Soc, 2008. 130(18): p. 5878–9.

51. Hussain, N.F., et al., Bilayer asymmetry influences integrin sequestering in raft-mimicking lipid mixtures. Biophys J, 2013. 104(10): p. 2212–21.

52. Vitrac, H., M. Bogdanov, and W. Dowhan, In vitro reconstitution of lipid-dependent dual topology and postassembly topological switching of a membrane protein. Proc Natl Acad Sci U S A, 2013. 110(23): p. 9338–43.

53. Mobashery, N., C. Nielsen, and O.S. Andersen, The conformational preference of gramicidin channels is a function of lipid bilayer thickness. FEBS Lett, 1997. 412(1): p. 15–20.

54. Lum, K., et al., Exchange of Gramicidin between Lipid Bilayers: Implications for the Mechanism of Channel Formation. Biophys J, 2017. 113(8): p. 1757–1767.

55. Nakano, M., et al., Flip-flop of phospholipids in vesicles: kinetic analysis with time-resolved small-angle neutron scattering. J Phys Chem B, 2009. 113(19): p. 6745–8.

56. Sapay, N., W.F.D. Bennett, and D.P. Tieleman, Thermodynamics of flip-flop and desorption for a systematic series of phosphatidylcholine lipids. Soft Matter, 2009. 5(17): p. 3295–3302.

57. Homan, R. and H.J. Pownall, Transbilayer diffusion of phospholipids: dependence on headgroup structure and acyl chain length. Biochim Biophys Acta, 1988. 938(2): p. 155-66.

58. Renooij, W. and L.M. Van Golde, The transposition of molecular classes of phosphatidylcholine across the rat erythrocyte membrane and their exchange between the red cell membrane and plasma lipoproteins. Biochim Biophys Acta, 1977. 470(3): p. 465–74.

59. Wolfenden, R. and M.J. Snider, The depth of chemical time and the power of enzymes as catalysts. Acc Chem Res, 2001. 34(12): p. 938–45.

60. Killian, J.A., K.N. Burger, and B. de Kruijff, Phase separation and hexagonal HII phase formation by gramicidins A, B and C in dioleoylphosphatidylcholine model membranes. A study on the role of the tryptophan residues. Biochim Biophys Acta, 1987. 897(2): p. 269–84.

61. Goforth, R.L., et al., Hydrophobic coupling of lipid bilayer energetics to channel function. J Gen Physiol, 2003. 121(5): p. 477–93.

62. Rokitskaya, T.I., E.A. Kotova, and Y.N. Antonenko, Tandem gramicidin channels crosslinked by streptavidin. J Gen Physiol, 2003. 121(5): p. 463–76.

63. Gurtovenko, A.A. and I. Vattulainen, Molecular mechanism for lipid flip-flops. J Phys Chem B, 2007. 111(48): p. 13554–9.

